# Both Prey and Predator Features Determine Predation Risk and Survival of Schooling Prey

**DOI:** 10.1101/2021.12.13.472101

**Authors:** Jolle W. Jolles, Matthew M. G. Sosna, Geoffrey P. F. Mazué, Colin R. Twomey, Joseph Bak-Coleman, Daniel I. Rubenstein, Iain D. Couzin

## Abstract

Predation is one of the main evolutionary drivers of social grouping. While it is well appreciated that predation risk is likely not shared equally among individuals within groups, its detailed quantification has remained difficult due to the speed of attacks and the highly-dynamic nature of collective prey response. Here, using high-resolution tracking of solitary predators (Northern pike) hunting schooling fish (golden shiners), we not only provide detailed insights into predator decision-making but show which key spatial and kinematic features of predator and prey influence individual’s risk to be targeted and survive attacks. Pike tended to stealthily approach the largest groups, and were often already inside the school when launching their attack, making prey in this frontal “strike zone” the most vulnerable to be targeted. From the prey’s perspective, those fish in central locations, but relatively far from, and less aligned with, neighbours, were most likely to be targeted. While the majority of attacks (70%) were successful, targeted individuals that did manage to avoid capture exhibited a higher maximum acceleration response just before the attack and were further away from the pike‘s head. Our results highlight the crucial interplay between predators’ attack strategy and response of prey in determining predation risk in mobile animal groups.

## Introduction

A key challenge in the life of most animals is to avoid being eaten. By living and moving together in groups, individuals can reduce their risk of predation (Ioannou et al., 2012; Krause and Ruxton, 2002; Pitcher and Parrish, 1993; Ward and Webster, 2016), benefiting from effects such as enhanced predator detection (Lima, 1995; Magurran et al., 1985), predator confusion (Landeau and Terborgh, 1986), and risk dilution effects (Foster and Treherne, 1981; Turner and Pitcher, 1986). The costs and benefits of grouping are, however, not shared equally. Besides differential food intake and costs of locomotion, individuals of the same group may experience widely varying risks of predation (Handegard et al., 2012; Krause, 1994; Krause and Ruxton, 2002). This not only has crucial implications for the selection of individual phenotypes, but also the emergence of collective behaviour, and the dynamics and functioning of animal groups (Farine et al., 2015; Jolles et al., 2020; Ward and Webster, 2016). Hence, better understanding what factors influence individual predation risk within animal groups is of fundamental importance.

Long-standing theoretical work predicts that, when predators appear at random and attack the nearest prey, predation risk should be highest on the edge (“marginal predation”) and front of mobile groups (Bumann et al., 1997; Hamilton, 1971;Morrell and Romey, 2008; Vine, 1971). Consequently if such predictions play out in the real world, individuals should try and surround themselves with others to reduce their domain of danger (Hamilton, 1971). There exists some empirical evidence of such “selfish herd” behaviour, in a range of species, such as individuals moving closer together when they perceive risk (Jakobsen and Johnsen, 1988; King et al., 2012; Krause, 1993; Sosna et al., 2019; Viscido and Wethey, 2002; but see Sankey et al., 2021). Several studies have also found evidence for predation risk being higher towards the edge (e.g. Krause, 1993; Romenskyy et al., 2020; Romey et al., 2008) and front (e.g. Bumann et al., 1997; Ioannou et al., 2019) of animal groups. However, while often assumed to be a universal pattern, the empirical evidence for this is equivocal and seems to be system-dependent, with a range of studies reporting that predation risk is actually higher in the group centre (e.g. Brunton, 1997; Hobson, 1963; Parrish, 1989) and towards the back of the group (e.g. Handegard et al., 2012; Krause et al., 2017). In relation to this, previous work has shown that individuals in such positions actually have poorer access to salient social information, as well as visual information of what happens outside the group (Rosenthal et al., 2015).

There has been a lot of focus in the literature on the spatial effects underlying predation risk, especially in terms of centre-to-edge and front-to-back positioning, partly because they are the easiest to measure, and research largely concentrating on a single or a few key features (but see e.g. Lambert et al., 2021; Romenskyy et al., 2020). However, a wide range of factors may be expected to influence predation risk. In particular, following Hamilton’s “selfish herd” theory (Hamilton, 1971), the spacing of individuals with respect to nearby neighbours, such as individuals with fewer neighbours/a larger domain of danger experiencing higher predation risk (De Vos and O’Riain, 2010; Lambert et al., 2021; Quinn and Cresswell, 2006; Romenskyy et al., 2020). But also individuals’ alignment to nearby neighbours (Ioannou et al., 2012), and features related to their visual information, which is known to strongly influence how well individuals respond to others (Davidson et al., 2021; Rosenthal et al., 2015; Sosna et al., 2019; Strandburg-Peshkin et al., 2013).

The majority of previous research has also focused exclusively on prey behaviour, building on the assumption of Hamilton’s classical model that a predator will attack the nearest prey (Hamilton, 1971). Thereby the predator is treated as an abstract source of risk, and hence any predator-related features as well the interactions among the predator and its prey are typically not explicitly evaluated. This is problematic as the behaviour and attack strategy of predators - such as to ambush, stealthily approach, or to hunt groups of prey - are likely to strongly influence which prey are ultimately attacked (Hirsch and Morrell, 2011; Lima, 2002; Stankowich, 2003), and will typically act as a strong selective force in the evolution of prey and predator traits (Lima, 2002). Furthermore, differential likelihood to be attacked is often used as a proxy for predation risk (Krause, 1994) because of difficulties in quantifying successful attacks (e.g. Handegard et al., 2012), predators failing to ever successfully attack (e.g. Parrish, 1989; Romenskyy et al., 2020), or it being impossible for predators to consume their prey (e.g. Ioannou et al., 2019; Milinski, 1977). Hence, there remains limited knowledge about the features that determine the probability that targeted prey may actually survive a predator attack.

To advance our understanding of differential predation risk in animal groups, and thereby of the evolutionary drivers of collective behaviour, we need to systematically investigate the relative influence of different group spatial, spacing, and visual features of both prey and predator and thus consider the real-time dynamics between live predators attacking groups of prey they can actually capture. This, however, poses a considerable challenge since it requires the simultaneous tracking of all members of a group of prey, as well as the predator, at a sufficiently high spatial and temporal scale. Here we present experiments in which we achieve this. Specifically, we observed Northern pike (*Esox lucius*), a geographically-widespread and ecologically-important predator (Craig, 2008), attacking free-swimming schools of 40 golden shiner fish (*Notemigonus crysoleucas*) to gain a detailed mechanistic understanding of when and where the predator attacks and what predict individual risk and survival.

We exposed pseudo-randomly composed groups (controlling for pike exposure) of juvenile shiners (8.5 cm, 95% CI: 6.4-11.5 cm) to individual pike (*n* = 13; 22.4 cm ± 1.1 cm) in a large open arena (1.05 m x 1.98m; 7 cm depth), and used custom-developed tracking software to acquire detailed spatial and temporal (120 fps) data for a total of 125 attacks (see Materials and Methods). By tracking both the predator and all prey individually over time, we were able to quantify each fish’s spatial position, relative spacing, orientation, and visual field, and analyzed their detailed movement kinematics throughout each attack (Figure 1). We then quantified how, when, and where the pike attacked and, subsequently, used model fitting procedures to infer the relative importance of a wide range of potential features to predict shiners’ risk to be targeted and their likelihood of surviving a predator attack using both a prey-focused and a predator-focused approach (see Table 1).

**Figure 1.**
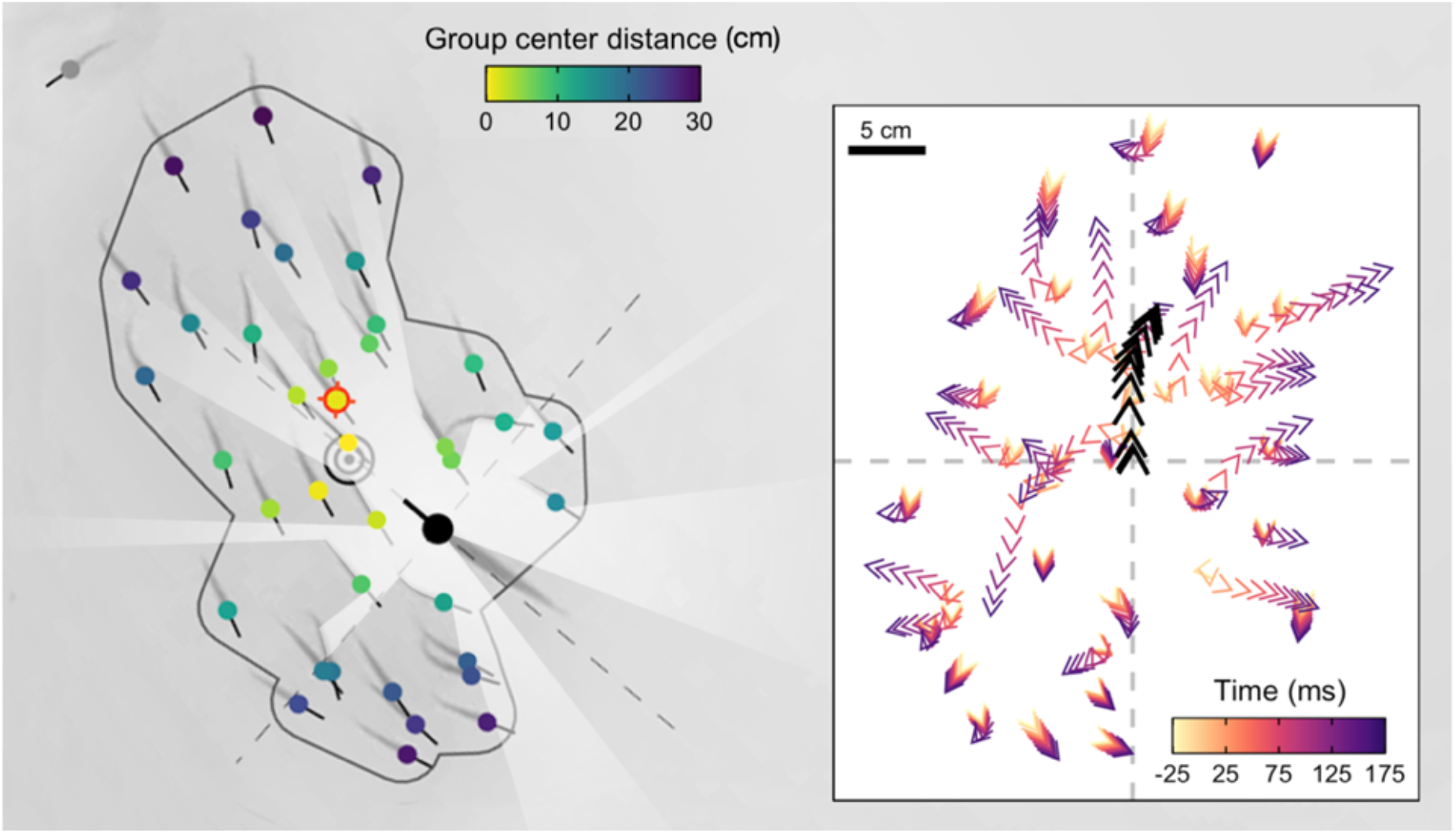
High-resolution Tracking of Predator Attacks. Cropped image from a sample video trial moments before the attack with key tracking data overlaid. Shiners are coloured yellow to blue based on their distance from the group centroid. Red target indicates the targeted individual, black concentric circles the group centroid, and the dark grey line the automatically determined school boundary, based on hierarchical clustering. Rays (white) represent a visualization of the pike’s field of view. Inlay figure presents detailed temporal data of the attack relative to strike initiation, with shiners positioned relative to the pike (black arrows) at the origin facing up.

**Table 1.** Description of features used in our multi-model inference approach for predicting which individuals were targeted and what explained their survival. For a visualisation of the features, see Figure 3.

| Feature | Acronym | Description |
| --- | --- | --- |
| Body length | BL | Shiner's body length (cm) |
| Centre distance | CD | Shiner's distance from the group centroid (cm) |
| Centre-edge position | CDrank | Shiner's CD ranked and scaled from 0 (most central) to 1 (least central) |
| Convex hull position | Hpos | Whether a shiner was part of the group hull or not |
| Inter-individual distance | IID | Shiner's median distance to all its group mates |
| Local misalignment | LMis | Difference in orientation angle (in degrees) between the shiner and its group mates within 10 cm |
| Voronoi area | VA | Area (cm<sup>2</sup>) around a shiner closest to that individual and not another individual, limited to the boundaries of the testing arena (log-transformed) |
| Limited domain of danger | LDOD | VA limited to a max radius of 10 cm from the shiner (log-transformed) |
| Front-back centre distance | FBCD | Shiners' distance from the group centroid in the plane of the group average orientation (positive values indicates in front of the centroid) |
| Front-back position | FBrank | Shiners' FBCD ranked and scaled from 1 (front) to 0 (back) |
| Visual weighted degree | WDeg | The proportion of each shiner's vision occupied by conspecifics |
| Distance to the pike | PD | Shiner's distance to the head centroid of the pike (cm) |
| Angle to the pike | PA | Shiner's position relative to the pike facing north (degrees), 0° being straight in front and 180° directly behind |
| Orientation to the pike | PO | Shiner's orientation relative to that of the pike |
| Pike vision of shiner | PVS | Pike's field of view occupied by the individual shiner (deg) |
| Target max speed | TMS | Targeted shiner's maximum speed in the 0.5s until the time of attack (cm/s) |
| Target max acceleration | TMA | Targeted shiner's maximum acceleration in the 0.5s until the time of attack (m/s<sup>2</sup>) |
| Target max turn | TMT | Targeted shiner's maximum orientation change (deg) in the 0.5s until the time of attack |
| Pike max acceleration | PMA | Pike's maximum acceleration in the 0.5s until the time of attack (m/s<sup>2</sup>) |

## Results

### How do pike attack groups of prey?

The pikes’ predatory movements typically began with an orientation phase in which they slowly turned their long body axis towards the schooling prey, followed by a stealthy approach (69% of attacks the pike moved steadily at < 0.5 BL/s), during which on average only 5% of shiners turned away (> 90°). After getting into position, the pike adopted an S-shaped body posture (see Appendix 1 Figure 1) to get ready for the actual attack – the strike – one sharp, sudden burst of movement (Figure 2A). By curving their body, the pike were able to generate a very rapid increase in kinetic energy (see further Domenici and Blake, 1997; Webb and Skadsen, 1980) and within a couple milliseconds attain a forward acceleration of 26.7 ± 0.7 m/s^2^ (mean ± SE), reaching speeds of 122.7 ± 3.6 cm/s, almost 1.5x higher than the escape speed of the prey they targeted (84.1 ± 2.7 cm/s; *χ^2^* = 104.7, *p* < 0.001; Figure 2B; reported values based on smoothed data). Due to its abrupt nature, we could automatically determine the exact moment of strike initiation at <0.01 s resolution, defined as the “time of attack” (see Appendix 1), and investigate the individual and collective behaviour of the prey and predator at this exact moment of the pikes’ predatory behaviour.

**Figure 2.**
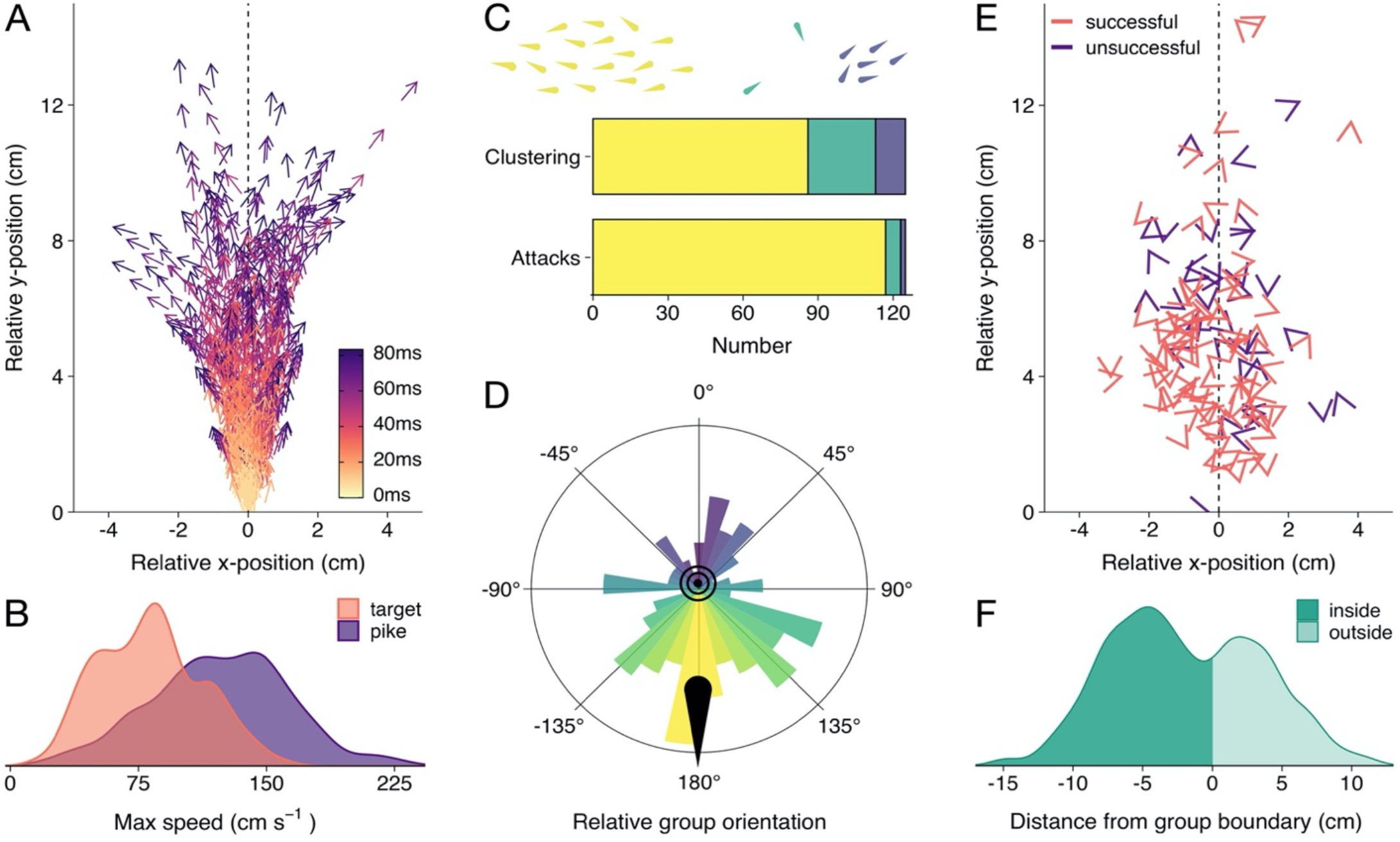
Detailed Attack Characteristics. (A) Pike attack trajectories that successfully resulted in prey capture (*n* = 88). Data are shown from the moment of time of attack (strike initiation), with the predator positioned at the origin facing north. (B) Density plots of the maximum (smoothed) speed of the pike and targeted shiners during the attack (from - 0.5s to +0.1s relative to strike initiation). (C) Barplots of shiners’ clustering (top) and pikes’ likelihood to attack different clusters (bottom). Top bar shows if prey were found in a single cluster (yellow), a single large cluster with small clusters of one or two individuals (green), or in multiple larger clusters (blue), as indicated by schematic above, while the bottom bar shows the number of attacks for each type of cluster. (D) Polar plot showing the distribution of group orientations relative to the pike facing north, coloured blue (0°) to yellow (−/+180°). (E) Positioning of targeted prey relative to the pike, with arrow headings indicating prey orientations. (F) Histogram of pikes’ distance from the group boundary. For figure D-F, data were subsetted to attacks of main cluster (*n* = 117) and focus on the time of attack.

### What is the collective state of the prey at the time of attack?

First, we determined if the shiners were generally found in one large school or multiple smaller groups using a hierarchical clustering approach. In short, fish were automatically clustered based on their inter-individual distance. If a large discontinuity in cluster distances was found, we considered there to be multiple groups of prey, based on a predetermined threshold (see Materials and Methods). However, the shiners were typically organised in one large, cohesive school at the time of attack, only occasionally with one or two singletons, and seldomly in multiple clusters of more than two fish (Figure 2C). In cases of attacks when there were multiple clusters, pike almost always (32/39 attacks) targeted an individual in the largest cluster (Figure 2C). We therefore focused all subsequent analyses on the 94% of attacks (*n* = 117) where the pike targeted an individual in the largest/only cluster, i.e. the main school.

In terms of the collective behaviour of the prey at the time of attack, the shiners were generally highly cohesive (mean inter-individual distance: 16.3 ± 0.4 cm), moderately well aligned (median polarization: 0.59), and moved at a modest speed (7.7 ± 0.6 cm/s, based on the group centroid). Rotational milling was rarely observed (mean group rotation order: 0.26), a state that is more characteristic of larger shoal sizes, as observed in similarly-sized experimental arenas (Davidson et al., 2021; Tunstrøm et al., 2013).

### Where do pike attack schooling prey?

To quantify if pike had a tendency to approach and strike the schools from a certain direction, we computed the groups’ centroid position, orientation, and heading (i.e. movement angle) based on all the shiners in the group, and transposed these to be relative to the pike facing north (0°). This revealed that pike had a strong tendency to attack individuals by approaching the groups head-on, both in terms of the groups’ relative orientation (circular mean: 170.7°, Rayleigh’s test: mean vector = 0.28, *p* < 0.001; Figure 2D) and direction of motion (when moving at >1.5 cm/s, *n* = 102; circular mean: 150.6°, mean vector = 0.25, *p* = 0.002). On average, pike launched their attack at 11.5 ± 0.6 cm from the group centroid and only 5.3 ± 0.2 cm from the prey they targeted (Figure 2E; 5.46 ± 0.3 cm including attacks where the pike did not attack the main cluster). While the shiners did not show a change in their packing fraction (median nearest-neighbour distance) with repeated exposure to the pike (*F*_1,52_ = 1.81, p = 0.185), they increasingly avoided the area directly in front of the pike’s head (Appendix 2 Figure 1A) resulting in the pike attacking from increasingly further away (target distance: *F*_1,52_ = 45.52, p < 0.001, see Appendix 2 Figure 1B,C).

Ranking individuals front to back (and scaling 1 to 0), we found that, rather than attacking individuals in the front of the group (see e.g. Bumann et al., 1997; Ioannou et al., 2019), pike tended to target individuals in relatively central positions (mean position: 0.45 ± 0.02; *n* = 117). Excluding groups with low polarization (< 0.4; n = 80), where it is more difficult to determine the “front”, did not substantially change this effect (0.48 ± 0.03). To investigate further if pike potentially launched some of their attacks from inside the school, we computed the smallest convex polygon that encompassed all individuals in the group and used concave approximations to create a realistic approximation of the group boundaries (see Figure 1). This revealed that indeed for more than half the attacks (63.2%), the pike was already partly inside the group boundary at the moment of attack (based on the location of the head centroid; Figure 2F). This observation could both be explained by the pike actively entering the group as well as the group moving to and around the pike. We therefore further computed the groups’ relative motion to that of the pike (see Materials and Methods), which revealed that for almost all attacks (92.3%) the pike moved more towards the school than the school moved towards the pike.

### What prey and predator features influence the predation risk of schooling prey?

To infer which features where the most predictive of individual predation risk of the schooling shiners, we used a multi-model inference approach (for details, see Appendix 3). This is a commonly-used technique whereby, rather than fitting a single model, models are fitted for every possible combination of features and their support is ranked based on information criteria (Grueber et al., 2011; Harrison et al., 2018). Features’ importance can then be assessed based on their relative weight across all models, with top-performing models being given more weight (Burnham and Anderson, 2002; Johnson and Omland, 2004). In our models we considered a combination of features where we had strong biological reasoning to be of potential influence (c.f. Burnham and Anderson, 2002), including those related to the spatial positioning, orientation, spacing, and visual field of both predator and prey (see Table 1 and Figure 3). Where relevant, ranked predictors, which place more attention on the relative differences between individuals, were also considered. We will first discuss the results from the perspective of the prey, and then from that of the predator.

**Figure 3.**
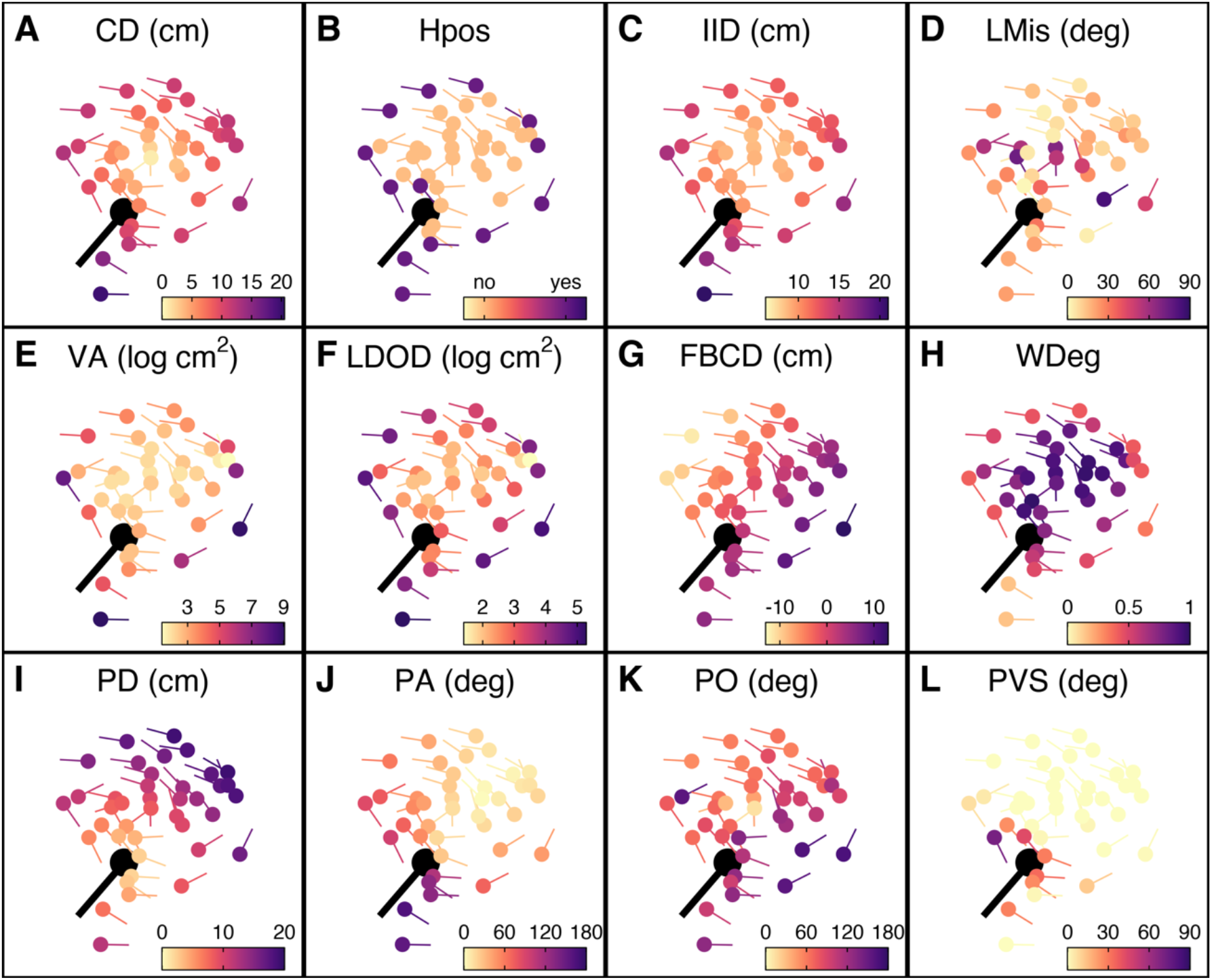
Prey features considered in the multi-model inference approach. (A-L) Schooling shiners and pike (black) at the time of attack with the shiners coloured based on the range of features used in our multi-model inference approach to predict which individual is targeted for attack. Visualisations show the positional and orientation data for a random representative trial and frame. An explanation of the acronyms can be found in Table 1. Note that ranked centre distance (CDrank) and front-back positioning (FBrank) are not included here but will visually resemble plots A and G. Also, for this particular example group rotation was high (0.81) and thus front-back positioning is not meaningful.

#### Prey-focused approach

For the “prey-focused approach” we ran multi-model inference using logistic regression considering all individuals in the school as potential prey and excluded any features about the predator (for further details about the models, see Appendix 3). Of the 11 features considered, three key predictive features emerged, as revealed both when considering feature weights (Figure 4A) and based on the top model (Appendix 3 – Figure 1A): (i) shiner’s ranked centre-to-edge position, (ii) shiner’s misalignment to surrounding neighbours (within a 10 cm radius), and (iii) the size of shiner’s limited domain of danger, the area around a shiner closest to that individual and not another individual, limited to a radius containing on average 25% of the other group members. In contrast to the widely-held assumption that predation risk is lowest in the group centre, we found that prey near the centre were more than twice as likely to be targeted than those near the edge of the group (scaled rank 0-1; estimate: −1.69 ± 0.37; LRT: *χ^2^* = 20.38, *p* < 0.001; Figure 4B). Multi-model inference also revealed that individuals were less likely to be attacked when they showed better alignment with their neighbours (estimate: 0.014 ± 0.004 misalignment in degrees; LRT: *χ^2^* = 15.01, *p* < 0.001; Figure 4D) and had a smaller limited domain of danger (LDOD), i.e. were surrounded closely by other groupmates (log area estimate: 0.41 ± 0.12; LRT: *χ^2^* = 12.99, *p* < 0.001; Figure 4C). Although LDOD was inherently smaller for prey the closer they were to the group centre (correlation coefficient: *r* = 0.52), there was considerable unexplained variance between these two features (R^2^ = 0.28). Prey’s front-back position, weighted degree (proportion of vision occupied by conspecifics), or whether they were positioned on the group edge or not had a much weaker influence on predation risk (i.e. the features were lower ranked, Figure 4A). Excluding groups with low polarization did not change the effect of front-back positioning (see Appendix 3).

**Figure 4.**
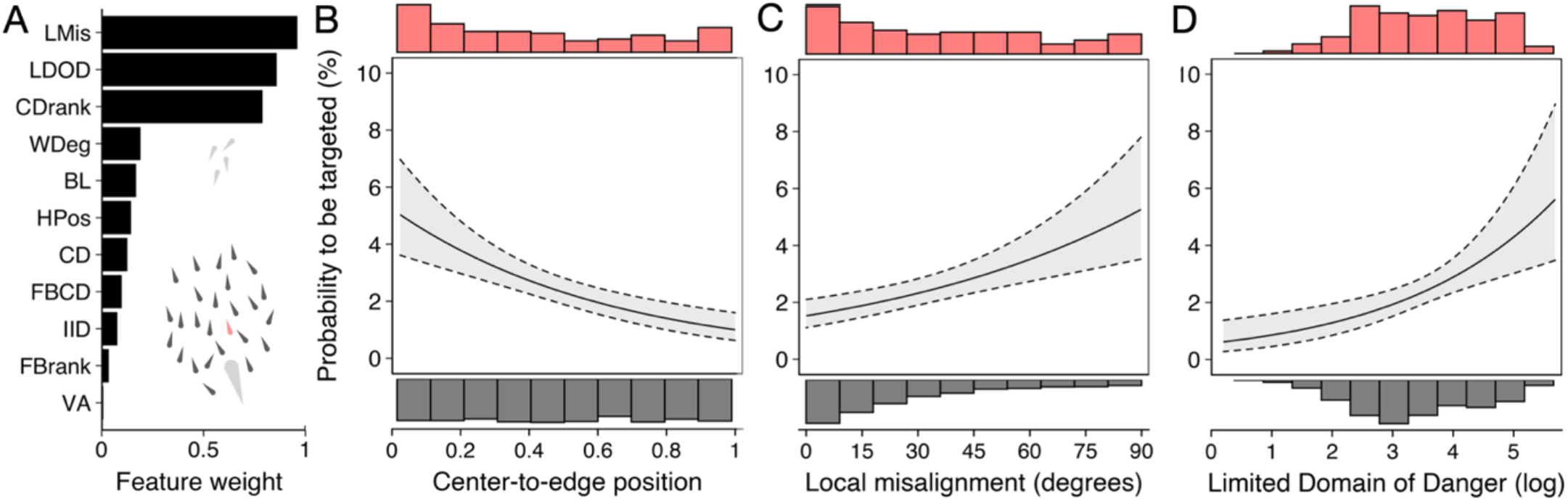
Predictors of the likelihood to be targeted - prey-focused approach. (A) Relative feature weights based on multi-model inference ranked from highest (top) to lowest (bottom) (for acronyms, see Table 1). (B-D) Top 3 features affecting the probability that an individual is targeted: (B) its ranked centre-to-edge position, (C) its misalignment with nearby neighbours (within 10 cm), and (D) its limited domain of danger (log-transformed). Plots are created using predicted values from the final model (see Appendix 3 Figure 1), with the envelope showing the 95% confidence intervals. Red and grey histograms are of the raw data of the targeted and non-targeted individuals respectively.

#### Predator-focused approach

By studying predation risk from the prey’s perspective, one ignores potentially crucial information about the predator’s attack strategy and decision-making (Hirsch and Morrell, 2011; Lima, 2002; Stankowich, 2003). To account for this, we reran multi-model inference but now also considered features about the predator, including shiners’ distance, angle, and relative orientation to the pike, as well as the proportion of pike’s vision occupied by each shiner (see e.g. Figure 3L). Furthermore, as most predators only have a specific region that they are biologically capable of attacking, we only considered shiners found inside the pikes’ strike zone, an area of roughly 8 cm wide and 15 cm long directly in front of the pike within which all targeted prey were positioned (Figure 2E). Using this predator-focused approach, we found as most predictive features (Figure 5A): (i) prey’s distance to the (head of) the pike (−0.514 ± 0.054; LRT: *χ^2^* = 146.82, *p* < 0.001; Figure 5B), (ii) prey’s angle to the pike (−0.070 ± 0.011; LRT: *χ^2^* = 61.32, *p* < 0.001; Figure 5C), and, as for the prey-focused approach, (iii) their limited domain of danger (0.675 ± 0.137; LRT: *χ^2^* = 26.02, *p* < 0.001; Figure 5D). Shiners were 6 times as likely to be targeted when they were within 6 cm and directly in front of the pike (−/+ 45°) compared to when positioned further away or more towards the side (49.1% vs 8.2% of attacks). To investigate if pike were likely to generally attack prey near to them, or the actual closest prey, one of the assumptions of Hamilton’s model (Hamilton, 1971), we ranked individuals based on their distance from the pike, again considering only individuals within the domain of danger. This revealed that for 73% of attacks (86/117) the pike did attack the nearest individual ahead, which increased to 90% considering the three nearest individuals. Consequently, swapping the feature of absolute pike distance with ranked pike distance considerably increased the predictive power of the model (ΔBIC = −37.4), thereby resulting in LDOD to drop as a significant feature.

**Figure 5.**
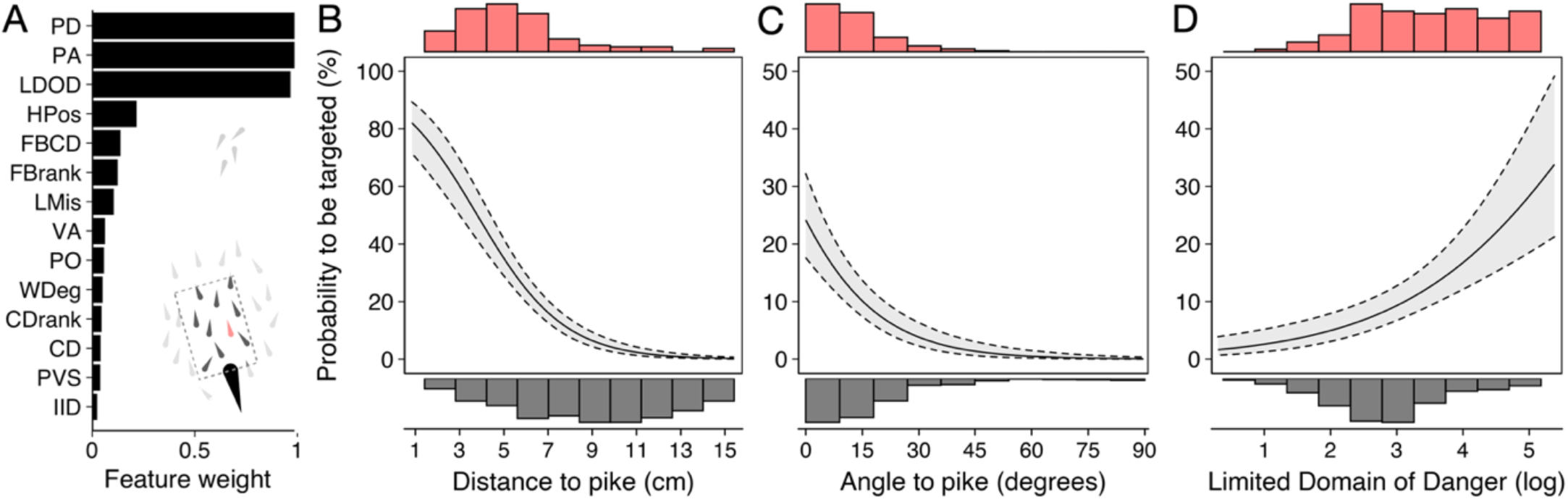
Predictors of likelihood to be targeted - predator-focused approach. (A) Relative feature weights based on multi-model inference and ranked from highest (top) to lowest (bottom). (B-D) Top 3 features affecting the probability that an individual is targeted using a predator-focused approach: (B) its distance to the pike, (C) its angle relative to the pike’s orientation, and (D) its limited domain of danger. Plots are created using predicted values from the final model (see Appendix 3 Figure 1), with the envelope showing 95% confidence intervals. Red and grey histograms are of the raw data of the targeted and non-targeted individuals respectively.

### Which features best predict predation avoidance success?

The pike in our study were relatively successful, with 70% of attacks resulting in the capture of prey. In contrast to previous work that only investigated the likelihood for an individual to be targeted (e.g. Handegard et al., 2012; Ioannou et al., 2012; Romenskyy et al., 2020), we could therefore also assess in detail what features are associated with targeted fish subsequently avoiding being caught. We compared individuals that were targeted yet successfully evaded capture (*n* = 34) with those individuals that were caught (*n* = 83), and considered as potential relevant features those that were found to be important in predicting which individual was targeted (see above), pike’s vision of its target at the time of attack, the targeted individual’s maximum speed, acceleration, and turning rate (max change in orientation), and pike’s maximum acceleration, all in the same standard 0.5 s time window until the time of attack. Running multimodel inference as before, two main predictive features emerged (Figure 6A; Appendix 3 Figure 2): (i) shiner’s maximum acceleration until the strike (Figure 6B) and (ii) shiner’s distance to the pike’s head (Figure 6C), which were themselves only very weakly related (4.9% of variance explained between them). Thus, targeted prey were more likely to evade capture when they showed a quick acceleration response in the moments before the pike launched its attack (−0.12 ± 0.04; LRT: *χ^2^* = 13.19, *p* < 0.001) and being further from its head (−0.27 ± 0.09; LRT: *χ^2^* = 8.98, *p* = 0.003).

**Figure 6.**
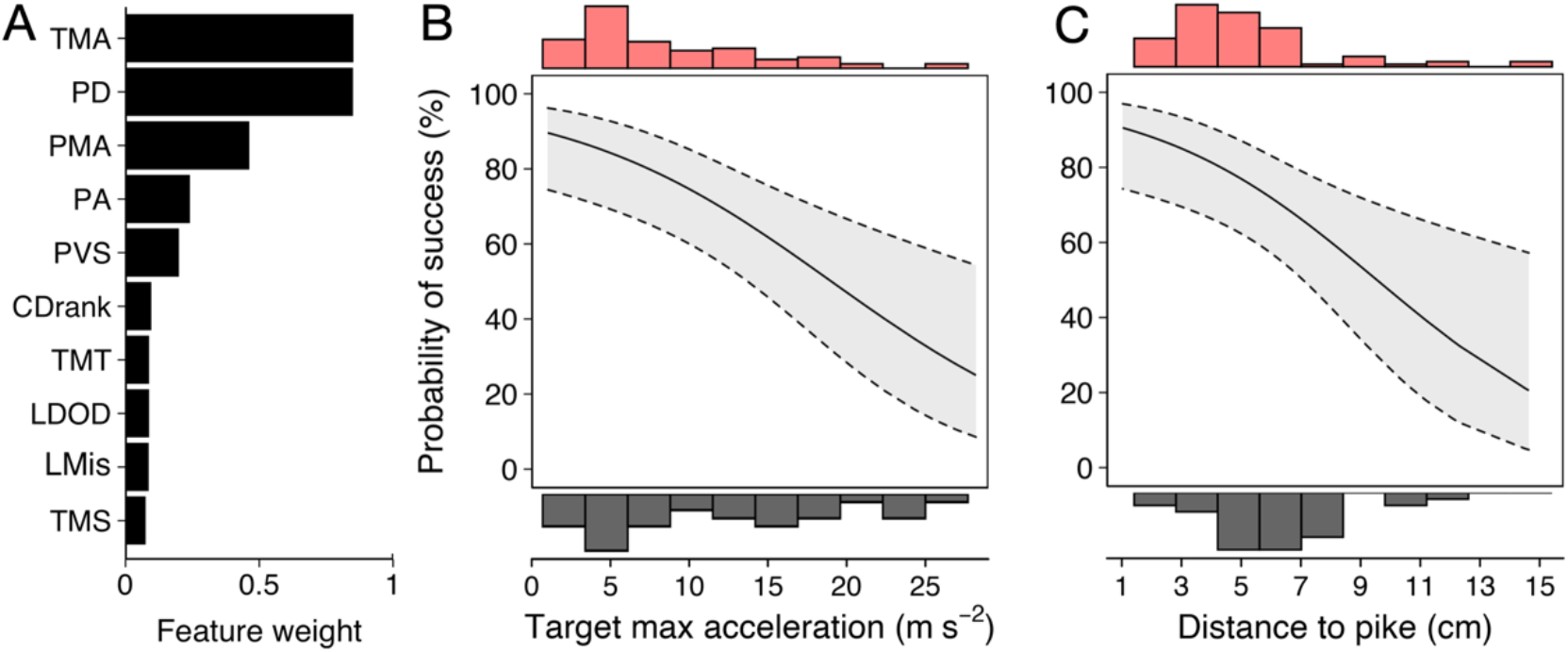
Predictors of Predator Attack Success. (A) Relative feature weights based on multi-model inference and ranked from highest (top) to lowest (bottom). (B-C) The two key features that predicted predator attack success: targeted shiners’ maximum acceleration in the half second before the attack (B), and their distance to the pike at the time of attack (C). Plots are created using predicted values from the final model, with the envelope showing 95% confidence intervals. Red and grey histograms are of raw data corresponding respectively to individuals that were targeted successfully and those that evaded the attack.

## Discussion

Understanding when, where, and how predators attack animal groups, and the types of anti-predator benefits grouping animals may experience, have been of long-standing interest (Ioannou et al., 2012; Krause and Ruxton, 2002; Ward and Webster, 2016). Although it is well appreciated that there is differential predation risk within animal groups, our understanding has, nonetheless, remained largely focused on marginal predation and selfish herd effects. By employing high-resolution tracking of predators attacking large, dynamically moving groups of prey, here we not only provide detailed insights into where and when predators attack, but also reveal the key features of predator and prey that influence which prey are targeted and what explains their chances for survival. Consideration of these multi-faceted factors underlying predation risk, in combination with predators’ attack strategy and decision-making, will have important consequences for understanding the costs and benefits of animal grouping and thereby the evolution of social and collective behaviour.

Pike tended to attack the groups head-on, in one fast burst of movement, following a slow and steady approach of the school. Rather than attacking individuals in the leading edge of the group, often assumed to be more risky than further back in mobile groups (e.g. Bumann et al., 1997; Ioannou et al., 2019; Krause et al., 2017; but see e.g. Handegard et al., 2012; Lambert et al., 2021), we found that pike, after steadily approaching the school, often slowly entered the school and then launched their rapid attack towards the group centre. That predators such as pike can get very close to their prey (see also Coble, 1973; Hoogland et al., 1956; Krause et al., 1998; Nursall, 1973; Webb and Skadsen, 1980) could potentially be explained by their narrow frontal profile (Webb, 1982), which makes it very hard for prey to detect movement changes, especially when attacked head-on. This may also explain why pike primarily targeted fish within the main school, even when stragglers or smaller groups were available, and the observation in previous work that pike may not suffer much from the confusion effect (Turesson and Brönmark, 2004).

Our finding that predators may actually enter groups during an attack and strike at central individuals is not often considered (Hirsch and Morrell, 2011), possibly because it contrasts with the long-standing idea that predation risk is higher on the edge of animal groups (Duffield and Ioannou, 2017; Krause, 1994; Krause and Ruxton, 2002; Stankowich, 2003). However, besides by predators using a stealthy approach, central individuals may also be more at risk by predators that attack groups from above (Brunton, 1997) or below (Clua and Grosvalet, 2001; Hobson, 1963; but see Romey et al., 2008), or that rush into the main body of a group (Handegard et al., 2012; Hobson, 1963; Parrish et al., 1989). Together, this suggests that central predation may be more widespread than currently considered and warrants broader research effort. In relation to this, the extent of marginal predation is actually predicted to depend on attack strategy and decline with the distance from which the predator attacks (Hirsch and Morrell, 2011), theory for which our study provides clear empirical support. Although internal positions being more risky in the context of attacks by ambush predators such as pike is likely not affected by group size, the relative risk of centrally positioned prey is expected to decrease with increasingly larger groups.

Previous work has often found that individuals on the edge and front of animal groups experience the highest predation risk (Bumann et al., 1997; Ioannou et al., 2019; Krause, 1993; Romenskyy et al., 2020; Romey et al., 2008). This supports the assumption of Hamilton’s classical model that the predator appears at random and strikes the nearest prey (Hamilton, 1971). Although here we also found that pike tended to attack the nearest prey (that they were physically capable of attacking), this prey was not actually the first they encountered. Pike tended to delay their attack until they were very close to and often even surrounded by their prey, clearly benefiting by being able to generate acceleration and speeding forces that were considerably higher than that of the prey they targeted (see further Domenici and Blake, 1997). Indeed, we found attacks were less likely to be successful when the target was further away and showed a high acceleration response in the moments before the strike. That pike very slowly and carefully approached the grouping prey, may thus be to minimize their strike distance, position and angle, until a point where they have to decide to either launch the attack or call it off based on the likelihood for a success (Nilsson and Eklöv, 2008). This type of predator decision-making *when* to attack, particularly relevant in the context of ambush predators, is rarely considered (Hirsch and Morrell, 2011), but fundamental for a proper understanding of the distribution of predation risk in animal groups across different predator-prey systems.

While many studies have investigated differential predation in animal groups, few have considered the many types of features that can be hypothesized to influence predation risk, let alone systematically compared them (but see e.g. Lambert et al., 2021; Romenskyy et al., 2020). Here, using a multi-model inference approach, we found that three key features stood out in influencing predation risk from the prey’s perspective: their centre-to-edge position, their limited domain of danger, and local misalignment. Individuals in the group centre were more at risk than those nearer the edge of the group. By closely approaching and attacking inside the group, pike may benefit by having more individuals to choose from within striking distance. However, in line with predictions from Hamilton’s selfish herd model (Hamilton, 1971), we also found that fish with a larger (limited) domain of danger were at higher risk of being targeted. Although some previous studies have shown individual predation risk is related to their domain of danger (De Vos and O’Riain, 2010; Lambert et al., 2021; Romenskyy et al., 2020), our results highlight its importance *within* groups, further supported by the finding of considerable unexplained variance between fish’s centre-to-edge position and their limited domain of danger. Such spatial heterogeneity within groups and its effects on differential predation is rarely considered (Jolles et al., 2020). That less aligned individuals had a considerably higher chance of being targeted is in line with experimental work using virtual prey (Ioannou et al., 2012) and could be a result of them simply stand out from their surrounding group mates (i.e. the oddity effect; Landeau and Terborgh, 1986) and/or because predators may have evolved to target misaligned individuals because of being less capable of obtaining salient social information (Couzin and Krause, 2003). Together, these findings bring some important nuance to the “selfish herd” phenomenon: rather than moving towards the group centre, individuals should try and position themselves near others, also inside the group and even when it is small or medium sized, and make sure they do not attract attention based on their orientation.

By rerunning multi-model inference with a predator-focused approach, i.e. by also including features about the predator and only considering prey within its perceived attack region, we found prey’s distance and angle to the pike were the strongest determinants for being targeted (see also Romenskyy et al., 2020). This shows that, besides moving towards others, as discussed above, it pays for prey to move away from the predator and avoid the cone of risk directly in front. By repeatedly testing the shiners with the pike, we saw that shiners indeed increasingly distanced themselves from the predator and especially avoided the region in front of the pike, consistent with previous work that suggests that, in addition to unlearned predispositions, experience with predators is important (Kelley and Magurran, 2003). While much previous work has only focused on predation risk from the prey’s perspective, these findings highlight that, for a proper understanding of predation risk in animal groups it is important to not remove the predator from the equation (Hirsch and Morrell, 2011; Lima, 2002). Interestingly, while vision is known to be the primary modality for pike’s predatory behaviour (Nilsson and Eklöv, 2008) and for mediating social interactions among species of schooling fish (Kotrschal et al., 1998; Rosenthal et al., 2015; Strandburg-Peshkin et al., 2013), we found that the extent that a shiner’s vision was occupied by conspecifics, its vision of the predator, and the predator’s vision of the shiner did not play a major role in determining predation risk.

Overall, the pike were successful, with 70% of attacks resulting in the shiner being eaten, which is comparable to previous studies with sit-and-wait predators (Krause et al., 1998; Neill and Cullen, 1974; Turesson and Brönmark, 2004; Webb and Skadsen, 1980). But where previous work was not able to quantify differential predation in terms of mortality risk, such as by using confined or virtual prey (Ioannou et al., 2019; Milinski, 1977), difficulties of field conditions (Handegard et al., 2012), or by the predator simply failing to ever attack successfully (Parrish, 1989; Romenskyy et al., 2020), here we were able to also quantify what factors determined the likelihood for individuals to survive an attack. We found that both predator distance and prey acceleration in the moments until the attack independently influenced predation success, with targeted prey that were further away, and that showed a faster acceleration response, being more likely to evade capture. These two features have also been found as significant variables affecting survival in a study on predatorprey dynamics with single prey (Walker et al., 2005; see also Lucas et al., 2021). Together, this suggests that targeted prey could sometimes anticipate the strike, highlighting that despite the very high speeds pike attained prey response does matter in predatorprey dynamics and that evasive prey behaviour may be especially successful when prey are further away (see also Ranåker et al., 2012). Although laboratory studies have their limitations, our approach has enabled us to acquire highly detailed individual-based characteristics of predator and all grouping prey as well as information about survival, thereby providing thorough new insights into predator behaviour and the predation risk in mobile animal groups.

In conclusion, using a quantitative empirical approach in which we acquired highly detailed individual-based characteristics of predator and grouping prey, we provide key mechanistic insights into when and where predators attack coordinated mobile groups and what predicts individual predation risk and survival. We demonstrate that ambush predators such as pike can stealthily approach grouping prey and delay their attack until being very close to, and often even until being surrounded by their prey. We also show that, rather than just go for the group centre, it pays for animals to position themselves near others, align with their body orientation, and avoid being positioned close to, and in front of, the predator, but that individuals still have a chance to escape an attack when targeted by avoiding the predator’s head and a strong acceleration response just before being attacked. Our study provides key insights about differential predation risk in groups of prey and highlights a fundamental role for both predator attack strategy and decision-making and prey behaviour. This is likely to have important repercussions for the costs and benefits associated with grouping, and ultimately the distribution of (social) phenotypes in populations (Farine et al., 2015; Jolles et al., 2020). It is therefore paramount for future work to consider the multi-faceted prey and predator features and the broader predation landscape that prey may encounter to properly understand of predator-prey interactions and the evolution of animal grouping.

## Acknowledgements

We thank the New Jersey Division of Fish and Wildlife for providing the pike used in this study and for their valuable advice, and Mike Gil for helpful feedback on a previous version of this manuscript. J.W.J. acknowledges support from a Alexander von Humboldt Postdoctoral Fellowship and a Postdoctoral fellowship from the Zukunftskolleg, Institute for Advanced Study at the University of Konstanz; M.M.G.S acknowledges support from a NSF-DDIG Graduate Research Fellowship (ID: 1701289); C.R.T acknowledges support from a MindCORE Postdoctoral Research Fellowship; J.B-C acknowledges support from the Center for an Informed Public and the John S and James L. Knight Foundation; I.D.C. acknowledges support from the Office of Naval Research grant (ONR, N00014-64019-1-2556), the European Union’s Horizon 2020 research and innovation programme under the Marie Skłodowska-Curie grant agreement (ID: 860949), the Struktur- und Innovationsfonds für die Forschung (SI-BW) of the State of Baden-Württemberg, the Deutsche Forschungsgemeinschaft (DFG, German Research Foundation) under Germany’s Excellence Strategy-EXC 2117-422037984, and the Max Planck Society.

## Author contributions

MMGS and JWJ conceived the ideas and designed the study with input from GPFM, JB-C, DIR, and IDC; MMGS performed the experiments; MMGS and JWJ processed the data, with tracking and validation software by CRT and JWJ, and visual field software by CRT; MMGS and JWJ ran initial data analysis; JWJ led the full data analysis and visualisations as well as the writing of the final manuscript, to which all authors contributed critically and approved its submission.

## Materials and methods

### Study species and animal holding

For our experiments we used golden shiners (*Notemigonus crysoleucas*) as prey and Northern pike (*Esox lucius*), a common predator of shiners (Johannes et al., 1989; Nursall, 1973), as predator. Shiners were purchased from Anderson Farms in Lonoke, Arkansas, USA and pike acquired from the New Jersey Division of Fish and Wildlife hatchery, where they were communally housed and fed a diet of fish pellets. After arriving at the Princeton University laboratories, fish were kept under controlled conditions (water temperature 16.5°C ± 1°C; room lighting: 12:12 hour light:dark cycle), with shiners and pike kept in separate rooms. Shiners were housed socially in 37-L glass tanks on a flow-through system with 20 individuals per tank and fed pelleted food (Zeigler Finfish) *ad libitum* once daily except for the day prior to an experimental trial. Pike were housed individually in 114-L tanks containing gravel and artificial plants, and were fed a single shiner every other day for the first 10 days after arrival after which they were only able to feed during the experimental trials (see details below). Experiments started after two weeks of acclimation in the laboratory. We ran two batches of experiments with approximately size-matched fish, with body size automatically determined with our tracking software (shiners batch 1: 7.9 cm, 95% CI: 6.7 - 9.8; shiners batch 2: 10.0 cm, 95% CI: 8.6 - 11.9 cm; pike batch 1: 20.0 ± 0.1 cm, *n* = 9; pike batch 2: 27.9 ± 0.4 cm, *n* = 4). The sex of the fish was not determined, but as shiners mainly school when they are juveniles, sex is not expected to play an important role. Animal care and experimental procedures were approved by the Princeton University Institutional Animal Care and Use Committee (protocol 2068-16).

### Experimental arena

Trials took place in a 3.5 × 6.5 ft (1.06 × 1.98 m) white Perspex tank. We designed the experimental arena to be large enough for the pike and shiners to move around freely and to facilitate free-schooling dynamics, based on the size of the fish and groups we were interested in. This was also informed based on previous experimental studies in which predators attacked schooling prey (e.g. Magurran and Pitcher, 1987; Neill and Cullen, 1974; Theodorakis, 1989). We expect that if a much larger space would have been used, pike would still show the same approach and attack behaviour linked to their stealthy attack strategy. External disturbances were minimised by placing the tank on two layers of carpet (to dampen vibrations), surrounding it by white curtains, and positioning it in an otherwise empty experimental room. Diffused light was provided by two LED panels positioned outside the curtains at the far ends of the tank. The tank was filled with water to a depth of 6 cm that had approximately the same temperature and water quality as that of fishes’ home tanks. The right-bottom corner of the tank contained an opaque Plexiglass partition to visually separate the pike from the shiners at the start of each trial. No refuge was provided during the experimental trials as pilots indicated that pike did not use them, in line with other experimental work with pike (Turesson and Brönmark, 2004). A Sony NEX-FS700 camera positioned at 2 m above the exact centre of the tank was used to film the experimental trials at 120 fps with a resolution of 1920 × 1080 pixels.

### Experimental procedure

An experimental trial started with netting the pike from the nearby holding room, transferring it to the experimental room in a covered bucket, and immediately releasing it into the partitioned corner of the experimental tank. While the pike was left to acclimate in the holding corner, a group of 40 shiners was taken from the separate holding room, transferred to the experimental room in a covered bucket, and released into the centre of the experimental tank. Subsequently, the experimenter closed the curtains around the tank, started video recording, and moved to a separate isolated section of the room. After five minutes, the partition was raised using a remote pulley system, giving the pike access to the whole tank. After another 10 minutes, recording was stopped and, to decrease potential stress, experimental lights were turned off and dim red lights turned on. One minute later, the shiners were carefully netted from the experimental tank and returned to the holding tanks and immediately fed. Subsequently, the pike was netted from the experimental arena in a gentle way and transferred to its home tank. After each trial, we drained and scrubbed the experimental tank thoroughly with a sponge to remove any potential predator and alarm cue odours.

A group size of 40 individuals was chosen as this reflects the size of shiner shoals observed in the wild (Hall et al., 1979; Krause et al., 2000), as well as that of many species that tend to form loose aggregations. For the experiments we aimed to get data for at least 100 attacks. As we also wanted to investigate how attacks potentially changed with repeated exposure, we could combine biological (different groups) and technical (independent repeated measures) replicates, thereby reduce the number of fish used in the experiments. We ran two batches of experimental trials with different shiners and pike. Three to four experimental trials were run per day, between 10:00 and 18:30, with three rest days between trials for each pike. Before the first trial, each pike was acclimated to the experimental arena by two mock trials with one and subsequently three shiners on two separate days, while each shiner group was allowed to acclimate to the experimental tank for 24 hours before their first pike exposure. To keep the experimental period consistent across all trials, trials continued for the full 10 minutes and were not stopped after the potential capture of a shiner. The order of testing was randomized and shiners were pseudo-randomly allocated to groups, accounting for repeated predator exposure. Specifically, after each trial, shiners were placed in tanks according to how often they had been tested already with the pike, and trials continued until there were not enough fish left for another trial with that number of exposures. Then a couple days later the next series of trials would start in the same way until all shiners had the same number of exposures again (up to a maximum of 6 exposures). Individuals were mixed across trials but making sure fish had the same number of exposures to the pike to avoid potential effects of familiarity between individuals and potential learning effects related to group composition or predator identity, as well as to always maintain a group size of 40. Pike had a mean number of 1.29 ± 0.08 attacks per trial, with 18 trials having more than 1 successful attack. Trials during which a pike did not attack were excluded and the pike and shiners were not used anymore for subsequent trials.

### Fish tracking

After automatically correcting all videos for minor camera lens distortion, we used custom developed tracking software to acquire highly detailed individual-based movement data for both the shiners (SchoolTracker, by Haishan Wu) and the pike (ATracker, by J. W. Jolles), including head position and body orientation. Full details of how SchoolTracker detects fish, tracks their movement, and corrects occlusions can be found in (Rosenthal et al., 2015); ATracker uses background subtraction algorithms and blob detection, shape, and threshold algorithms to get the pikes’ position and orientation while excluding the shiners. For our analyses we assured perfect tracks linked to each individual shiner’s identity in the large schools at 120 fps from one second before strike initiation through to one second later, and manually corrected the tracking data where needed. By tracking from above, we acquired detailed data on the predator attacks in two dimensions. We do not expect considerable differences when instead groups would have been tracked in 3D, foremost because schools of shiners tend to form flat schools near the water surface (Hall et al., 1979; Stone et al., 2016) but also because recent experiments have shown that adding the third dimension did not significantly improve the predictability of predation risk (Romenskyy et al., 2020).

### Quantification of behaviour

After tracking, we smoothed the positional coordinates using a Savitzky-Golay smoothing filter with a window of 0.1 s and converted pixels to mm. Using the head as reference, we then computed each fish’s velocity, speed, and acceleration as well as their distance to the closest wall. For each shiner we also computed their distance to the pike, their angle to the pike (absolute, with 0° being directly in front of and 180° directly behind the pike), and relative orientation to the pike (with 0° being the same orientation as the pike and 180° an opposite orientation). We did this by shifting the coordinates of the fish so that the origin of the coordinate system was at the pike and rotated such that the pike was facing up (0°) to have a common frame of reference. The maximum speed and acceleration values we report in the text are based on the smoothed data, which helps overcome the issue that already tiny shifts in reference points in subsequent frames during tracking can confound single data values. Computing these metrics on unsmoothed data instead results in maximum speed and acceleration values that are in line with that typically reported for attacks or escape (Domenici and Blake, 1997; Walker et al., 2005).

Next, we used a hierarchical clustering approach to determine the distribution of shiners in one or multiple groups. In short, we computed clusters hierarchically by starting with each shiner assigned to its own cluster and iteratively joined the two most similar clusters based on the centroids of those clusters until there was just a single cluster. The optimal number of clusters was determined automatically based on the change in distance between the clusters’ centroids in a single step relative to the variance in cluster distance. We thereby looked for large discontinuities in the change in cluster distance and used a predetermined threshold to optimize the clustering. This approach provides more realistic clustering than other approaches, such as those that use a fixed distance measure for clustering. For all attacks we made sure to manually check the computed clusters at the frame of attack, and in a few cases made a correction when there was a large disparity in body orientation (>90°) between a fish and the rest of the group. For each attack we then ranked the clusters and focused subsequent analyses on all attacks where pike attacked the largest cluster (*n* = 117). To get a realistic representation of the boundaries of the group (i.e. the largest cluster), we ran convex and subsequently concave hull approximations based on the position of all shiners in the group and scored if fish were on the group boundary as well as pikes’ distance to the group hull, with negative values indicates the pike was inside the group boundaries.

For each shiner we quantified their Voronoi polygon area, limited domain of danger (LDOD; following Lambert et al., 2021), local misalignment, and their inter-individual distance (see Table 1 for an explanation how these measures were computed). LDOD and local misalignment require a spatial threshold outside of which fish will be included. We considered the distance within which fish on average had 25% of their neighbours, which provides a good balance between generating enough variation among individuals and the area containing too many individuals. As across all trials fish on average had 25% of their neighbours within a distance of 9.9 cm, we used a threshold of 10 cm to include neighbouring fish for our LDOD and local misalignment measures. At the group level, at each time point throughout the attacks we determined the group’s (i.e. the largest cluster) position and movement vector based on its centre of mass and calculated group speed, cohesion, the average inter-individual distance between all shiners, polarization, which is a measure of the alignment of the fish in the group relative to each other and ranges from 0 (complete non-alignment) to 1 (complete alignment), and milling, which is a measure of high local but low global alignment as the group rotates around its core (Tunstrøm et al., 2013). Subsequently, we computed shiner’s absolute distance to the group centroid, and their distance along the front-to-back axis (positive when ahead of the centroid, negative when behind). As for the individual shiners, we computed the groups’ position, heading, and orientation relative to that of the pike. To make sure group relative heading changes were not due to changes in pike position, we calculated the change in position of the group relative to the pike from 0.1s before strike initiation while keeping the pike’s position constant. We thereby considered the same fish such that group centroid position could not fluctuate due to potential changes in cluster size. This movement of the group centroid equates to the average direction of motion of the prey fish.

Finally, to estimate the visual information available to each fish we used a ray-casting algorithm, originally developed for Rosenthal et al. (2015) (for further details see their paper). Visual features computed using this method have been shown to be informative of evasion behaviour (Rosenthal et al., 2015; Sosna et al., 2019), even in field conditions (Hein et al., 2018). We used the visual information to compute the proportion of each shiner’s vision that was occupied by conspecifics (weighted degree) and the proportion of a shiner that was visible to the pike. While individuals form a relatively planar group structure near the water surface, as schools are not perfectly 2-dimensional, it may be the case that neighbouring individuals do not always block an external view. However, shiners tend to form relatively planar groups near the water surface (Hall et al., 1979; Stone et al., 2016), and using incomplete rather than full blockage seems to have very little effect on an individual shiner’s detection coverage (Davidson et al., 2021).

### Analyses

To investigate if pike had a higher maximum speed than the shiner they targeted, we computed both fishes’ (smoothed) speed from 0.5 s before until 0.1 s after strike initiation and ran a linear mixed model with fish as fixed factor (predator; prey), maximum speed as response variable, and attack id as a random factor. To determine if groups were more likely to attack the groups head-on in terms of their orientation and direction of motion/heading, i.e. if their angles concentrated around 180° relative to the pike oriented north (0°), we ran Rayleigh tests for uniformity, with the Rayleigh statistic varying from R = 0, indicating a uniform distribution in all directions, to R = 1, representing that all vectors point in the same direction at 180°. For the analysis regarding the group heading, we made sure to subset the data to attacks where the group was moving at least at a speed of 1.5 cm/s (*n* = 102). To determine what features were predictive of whether a shiner was targeted by the pike and survives an attack, we used a multi-model inference approach (Burnham et al., 2011), focusing our analyses on all attacks where the pike attacked the main school (*n* = 117), which is described in much detail in Appendix 3.

## Appendices

### Appendix 1. Quantification of the time of attack

Pike attacks are characterized by a sharp, sudden, burst of movement, the “strike”. To automatically determine the exact frame of strike initiation (“time of attack”), we quantified the work rate by looking at the change in kinetic energy c.f. (Sosna et al., 2019). In short, for an individual *i*. with mass *m* and speed s, the change in kinetic energy is given by: Δ*KE_i* = *ms_i* * ((*ds_i*)/*dt*). As here we simply use kinetic energy to determine the frame of strike initiation, we considered the mass as a constant across all pike. We then rescaled this measure by 1/*m* to get “swimming intensity”. To automatically differentiate strikes from normal swimming, we focused on the peak intensity an individual attained and compared that to a swimming intensity threshold of two standard deviations above its median intensity in the second (120 frames) prior to its peak intensity during an attack. Normal swimming lacks a simultaneous high speed and acceleration, thus enabling us to automatically and objectively determine the exact time of attack at 120 fps (see Appendix 1 Figure 1), which was confirmed by manual video inspections.

**Appendix 1 Figure 1.**
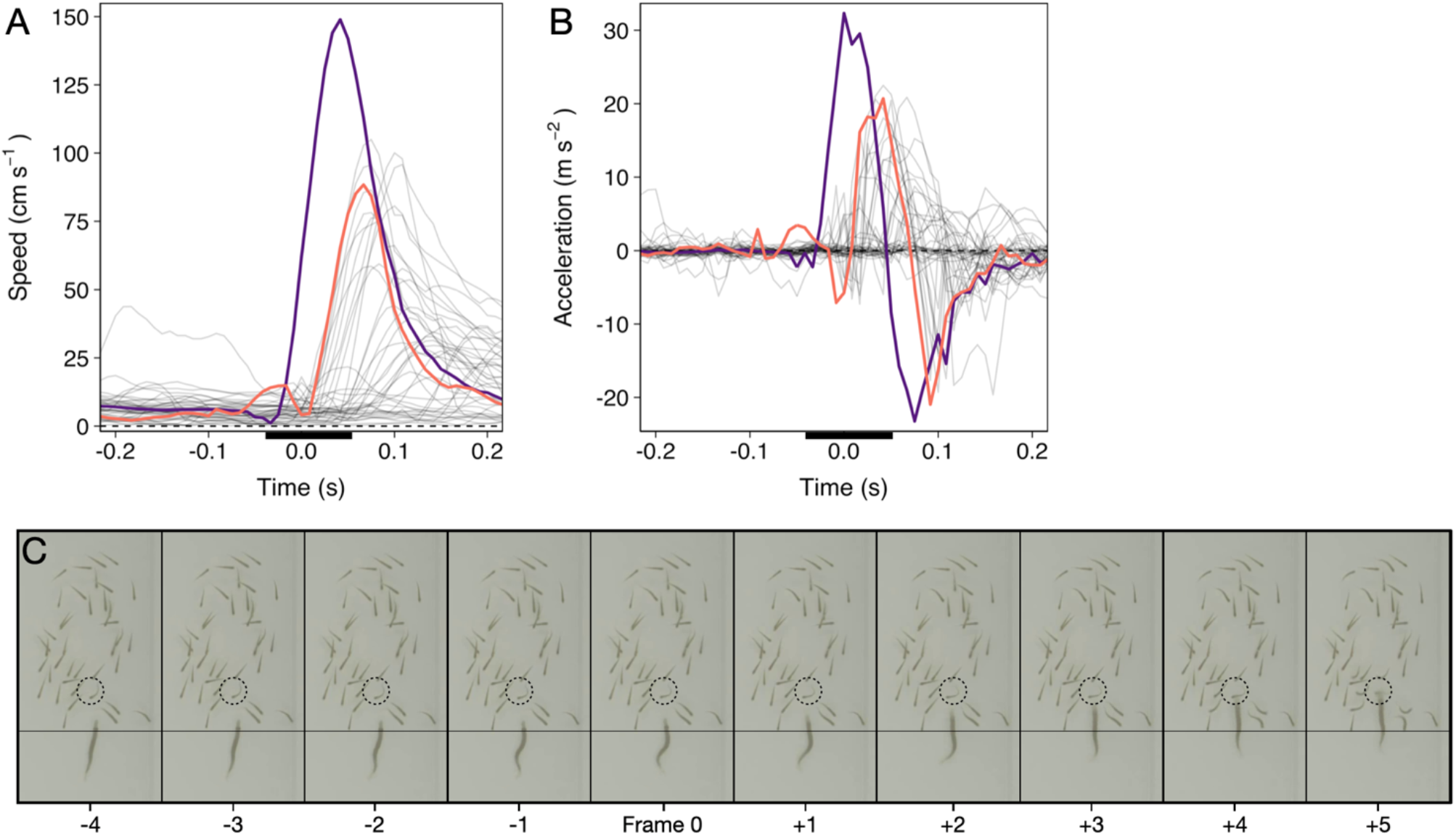
Time series of a randomly selected pike attack. (A and B) Speed and acceleration curves (based on smoothed data) of the pike (purple) and shiners (grey; targeted shiner in orange). Time is relative to the automatically quantified time of attack. Black bar reflects time series of panel C. (C) Cropped screenshots of the frames around the strike (indicated by the black bar in panels A and B), showing the characteristic S-shaped body posture leading up to the strike, during which the pike’s head stays roughly at the same position (see thin horizontal line). Targeted individual is indicated by the dashed circle. This specific strike lasted a total of 6 frames, or 0.05 s, until impact.

### Appendix 2. Effect of repeated exposure

**Appendix 2 Figure 1.**
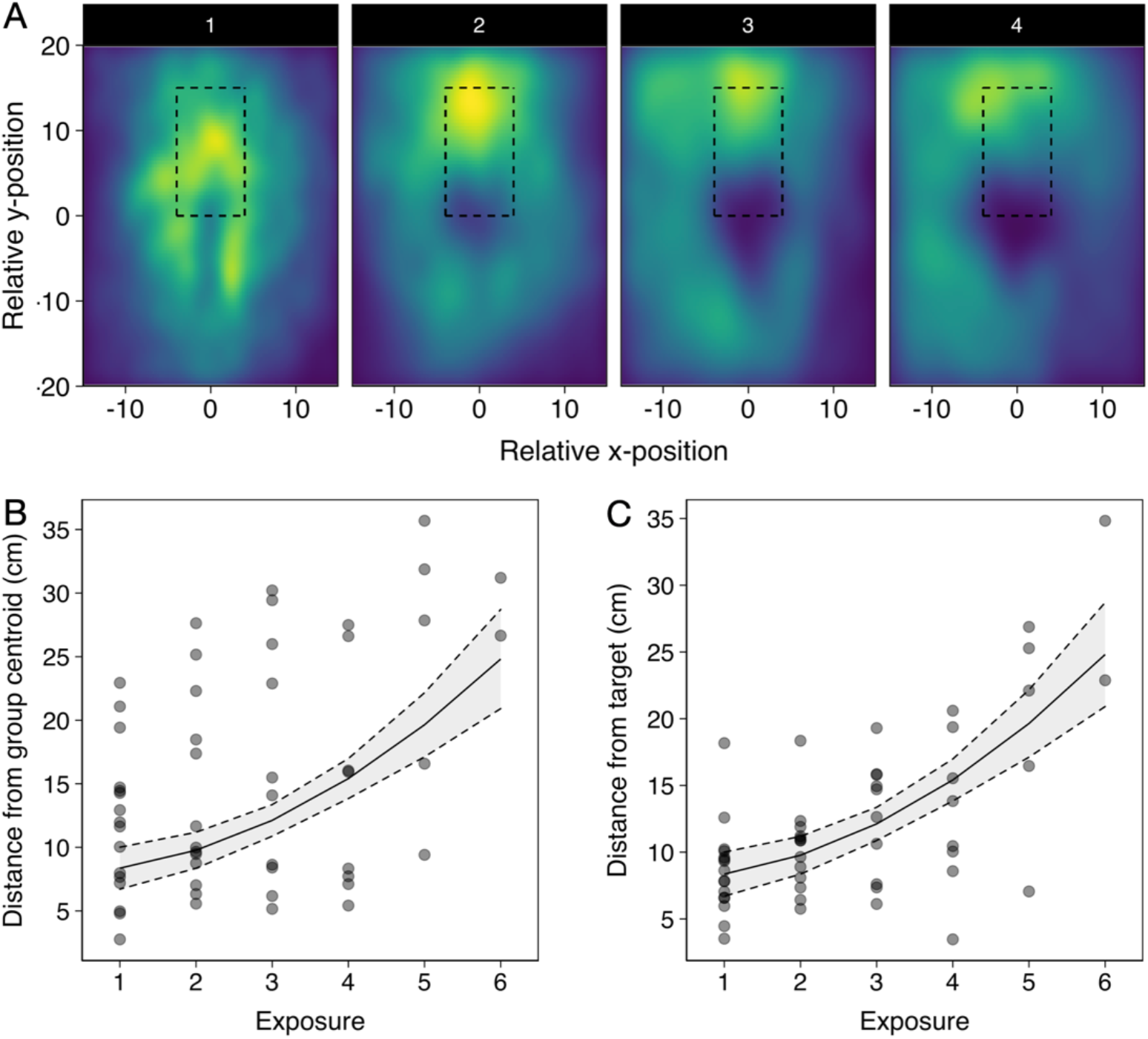
(A) Heatmaps of the relative position of all shiners to the pike up to strike for all first attempts relative to exposure, with the 4^th^-6^th^ exposure (panel 4) being grouped due to their smaller sample size, and data subsetted to the relevant region around the pike (nr of datapoints, i.e. a shiner location at a given frame, per attempt: 1 = 650.000; 2 = 528.950; 3 = 325.570; 4+ = 319.995). Rectangular area shows the area in which all attacks occurred. (B and C) Plots of the relationship between pikes’ distance to the group centroid (B) and to the targeted individual (C) at 0.5 s before the strike with repeated exposure. Line and 95% confidence intervals (dashed lines) are from a linear mixed model with exposure fitted as a cubic function. Data was subsetted to pikes’ first attack attempt during the trials.

### Appendix 3. Feature selection for predictors which shiners are targeted and survive attacks

To determine what features are predictive of whether a shiner i) is targeted by the pike and ii) survives an attack, we used multi-model inference (Burnham et al., 2011), focusing our analyses on all attacks where the pike attacked the main school (*n* = 117). In multi-model inference, rather than fitting a single model, a model is fit for every possible combination of features (predictor variables) and the relative importance of each feature can be assessed based on which features are present in the best models. We included a large range of potentially relevant features to characterise the prey and thereby determine their likelihood to be attacked (Appendix 3 Figure 1). We then ran exhaustive (i.e. not stepwise) model evaluation of all possible feature subsets to determine optimal feature importance (Johnson and Omland, 2004). Models were weighted based on their BIC score, which places a larger penalty on regression coefficients than AIC to avoid overfitting, after which relative feature importance was calculated based on all models that contained the feature, following (Burnham et al., 2011; Rosenthal et al., 2015). We thereby excluded any models that contained both absolute and ranked variables of the same feature (e.g. distance to centre and ranked centre-to-edge position) as these were inherently strongly correlated. We also made sure other included features did not show high collinearity by checking their Variance Inflation Factor (using a VIF criterium of 5, Harrison et al., 2018). The final predictive features were selected based on their relative importance score across all models as well as by considering the features that appeared in the most parsimonious models (see Appendix 3 – Figures 1 and 2).

**Appendix 3 Figure 1.**
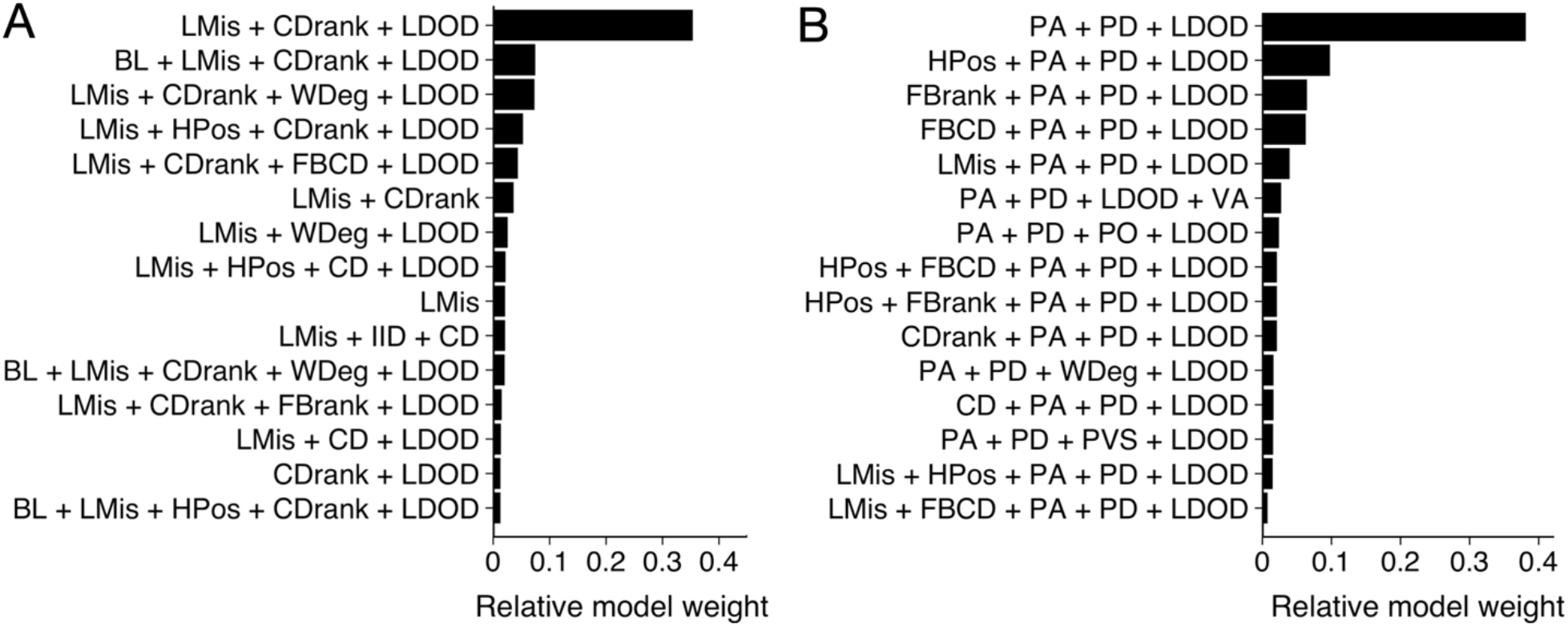
Model results from multi-model inference on likelihood to be targeted. Results from multi-model inference investigating which individual is targeted for attack using the prey-focused (A) and predator-focused (B) approach. Panels show relative evidence weight for the top 15 models for each approach. For acronyms, see Table 1 in the main text.

We started with a prey-focused approach to investigate predation risk purely from the prey’s perspective and thereby considered all shiners that were part of the main school to be potential targets for the pike. We ran generalized linear models with a logistic link function, with as response variable whether an individual was targeted or not (1,0). To account for the hierarchical structure of our data, we included as random effects exposure, Pike ID and attack attempt (1^st^, 2^nd^, 3^rd^) nested within trial. As potential predictive features we included distance to the group centroid (cm), front-back centroid distance (cm), Voronoi area, Voronoi area limited to a maximum radius of 10 cm from a conspecific (LDOD), median inter-individual distance (cm), weighted degree, and whether an individual was positioned on the group boundary (yes/no). We also included ranked variables for distance to group centroid and front-back centroid distance, further scaled between 0 and 1 to account for variation in cluster size, which better reflect individuals’ relative positioning in the group. To investigate predation risk from the predator’s perspective, we also considered features that included information about the pike, specifically shiners’ distance, angle, and relative orientation to the pike, and how much the pike saw of each shiner. Furthermore, we now only considered shiners within the attack region, the region in which all attacks occurred, 8 cm wide and 15 cm long directly in front of the pike. To quantify what best predicted an individual’s likelihood to survive an attack, we used predation success as our (binary) response variable and again ran multi-model inference, considering all significant features that arose from our prey- and predator-focused approaches looking at which individual was targeted. We additionally included pikes’ maximum acceleration in the 0.5 s prior to strike initiation, shiners’ maximum speed, acceleration, and turning angle in the 0.5 s prior to strike initiation, and pike’s vision of the shiner. As targeted shiners’ maximum speed and acceleration were positively linked (*r* = 0.68), all models that contained both these features were excluded. Furthermore, measures of maximum speed and acceleration throughout the full attack were not included as these could be confounded by the difference in duration of the attacks observed. How much the targeted shiner saw of the pike was not included either because this measure was found to be strongly linked to pike’s vision of the shiner (*r* = 0.40).

Our measures of local misalignment and LDOD were quantified for all neighbouring fish within a radius of 10 cm as this on average equated 25% of the school. To make sure the range over which we investigated local misalignment did not influence its predictive power, we also ran a model for the prey-focused approach with local alignment computed for individuals within a topological neighbourhood of 6 neighbours (c.f. Ballerini et al., 2008). Although this slightly weakened model fit (ΔBIC = +8.9) it did not qualitatively change the model results. Similarly, as front-back positioning is not meaningful for schools that are not well aligned, we re-ran model selection on the data subsetted to schools with a group polarization of at least 0.4 (*n* = 80 attacks). The resulting model output was not qualitatively different from the final model of feature selection, and adding front-back positioning as an additional feature again weakened model fit (ΔBIC = +6.3). Pike’s distance to the tank walls could also be hypothesized to influence its attack success. Adding this variable to the final model did however show this was not the case (ΔBIC = +4.4).

**Appendix 3 Figure 2.**
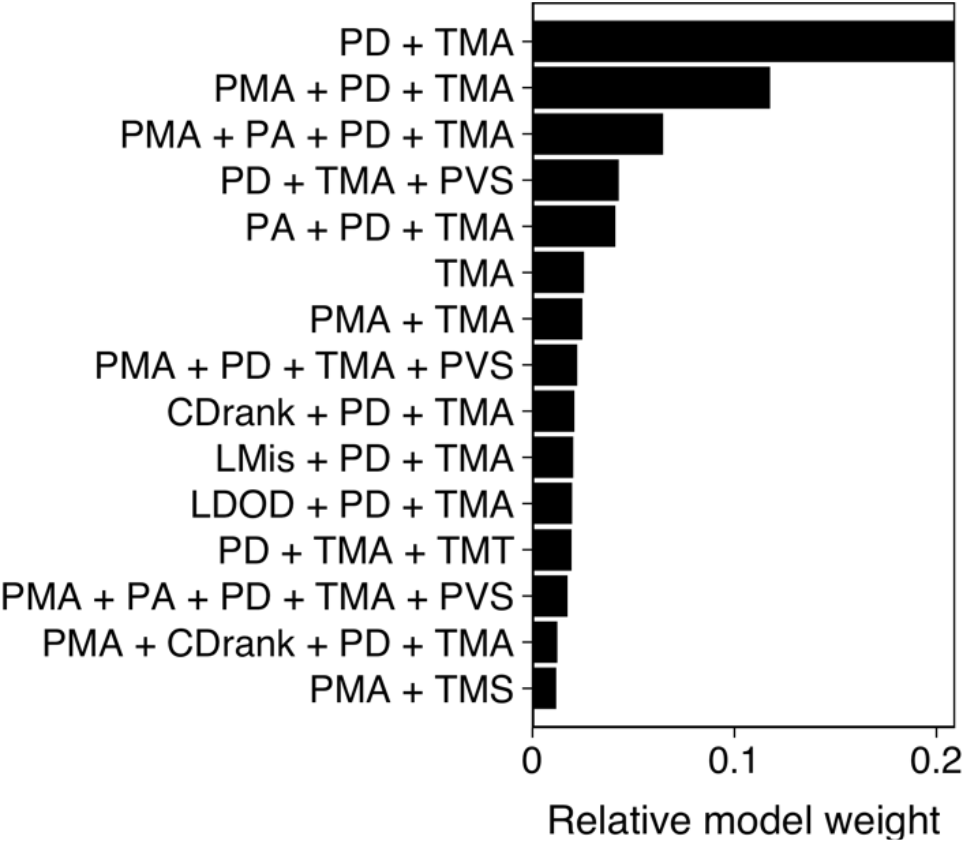
Model results from multi-model inference on predation attack success. Panel shows relative evidence weight for the top 15 models. For acronyms, see Table 1 in the main text.

